# Significant metabolic improvement by a water extract of olives: animal and human evidence

**DOI:** 10.1101/151860

**Authors:** Nikolaos Peroulis, Vasilis P. Androutsopoulos, George Notas, Stella Koinaki, Elsa Giakoumaki, Apostolos Spyros, Εfstathia Manolopoulou, Sophia Kargaki, Maria Tzardi, Eleni Moustou, Euripides G. Stephanou, Efstathia Bakogeorgou, Niki Malliaraki, Maria Niniraki, Christos Lionis, Elias Castanas, Marilena Kampa

**Author notes:** Corresponding Author All correspondence should be addressed to Dr Marilena Kampa,* Laboratory of Experimental Endocrinology, University of Crete, P.O. Box 2208, Heraklion, 71003 Greece.

## Abstract

Dyslipidemia and impaired glucose metabolism, are main health issues of growing prevalence and significant high Health Care cost, requiring novel prevention and/or therapeutic approaches. Epidemiological and animal studies revealed olive oil as an important dietary constituent for normolipidemia. However, no studies have specifically investigated the polyphenol rich water extract of olives (OLWPE), generated during olive oil production. Here, we explore OLPWE in animals and human metabolic parameters. High fat-fed rats developed a metabolic dysfunction, which was significantly impaired when treated with OLWPE, with decreased LDL and insulin levels and increased HDL. Moreover, they increased total plasma antioxidant capacity, while several phenolic compounds were detected in their blood. These findings were also verified in humans that consumed OLWPE daily for four weeks in a food matrix. Our data clearly show that OLWPE can improve glucose and lipid profile, indicating its possible use in the design of functional food and/or therapeutic interventions.

## INTRODUCTION

Impaired glucose tolerance and lipid metabolism are the most common metabolic dysfunctions in humans and they have been closely associated with obesity, now recognized as a chronic disease of alarming incidence (close to 40% of adults in the world are overweight or obese) ^1^. Obesity complications can further result to a number of life-threatening pathological conditions. In fact it is the various metabolic disorders (such as dyslipidemia and impaired glucose tolerance) are usually seen in central type obesity ^2^ together with increased blood pressure, that characterize a pro-inflammatory state ^3,4^, leading to an increased likelihood of insulin resistance/type 2 diabetes, atherosclerosis/cardiovascular disease ^5^. This, together with a resulting pre-thrombotic state ^6^ may result in premature death. This, obesity induced cascade of events characterizes a pathophysiological state, commonly referred as “metabolic inflammation”. This is a low-grade inflammation triggered by adipose tissue hypertrophy and hyperplasia and subsequent hypoxia. As a result, there is an altered lipid metabolism and an increased production of several hormones, chemokines and cytokines and coagulation factors that lead to hyperinsulinemia, β pancreatic cell dysfunction, type II diabetes, increased sodium uptake, vasoconstriction and hypertension, increased lipoprotein synthesis, gluconeogenesis and dyslipidemia, endothelial dysfunction, atherosclerosis and clotting disorders and ultimately to coronary heart disease. The collective cluster of (central) obesity, dyslipidemia, impaired glucose tolerance/insulin resistance and hypertension is commonly refered as the “metabolic syndrome”^7^

Obesity, due to its high global prevalence and comorbidities, has become an international health care priority, with the major aim being early diagnosis of metabolic dysfunction and improvement of body weight and adverse metabolic disturbances (mainly lipids and glucose) by dietary modifications and pharmaceutical interventions. The cost of the latter is extremely high ^8,9^and therefore alternative approaches, which may improve the above elements, are of great importance, both for the health of the patients and for a possible reduction of the pharmaceutical expenditure.

A great variety of animal and human epidemiological studies, report beneficial effects of olive oil and / or olive -olive oil polyphenol extracts ^10-12^ on glucose and lipid metabolism. However, during olive harvesting and olive oil production, a water phase is also produced, commonly discarded. This phase is rich in olive (poly)phenols, which distribute between the olive and water phase as a result of time of olive paste malaxation and temperature. Here, we studied the effect of a microencapsulated olives water phenolic extract (OLWPE) in a rat model of diet induced obesity and extended our study (as a proof of principle) in a healthy human population that consumed this extract in the context of a food matrix. Our findings clearly show that OLWPE can ameliorate main metabolic parameters, such as fasting glucose levels and lipid profile, indicating its possible dietary and/or therapeutic use.

## RESULTS

### Polyphenolic characterization of OLWPE

The phenolic content of concentrated OLWPE was initially estimated as 10mg/ml Trolox equivalents, while NMR analysis (Figure 1A), revealed that the extract, as expected, contains phenolic compounds along with other larger amounts of small molecular weight chemicals, such as ethanol, lactic, succinic and acetic acid and carbohydrates. Analysis of the phenolic area of the spectrum revealed that the main phenolic compound present is the phenylethanoid tyrosol, along with small amounts of ligstroside and possibly elenolic acid, which is a common phenylethanoid hydrolysis product. Further analysis of the extract by the more sensitive LC-ESI-MS/MS method showed also the presence of the phenylethanoids oleuropein and verbascoside, the flavanols catechol, catechin, and epicatechin, the flavones apigenin, apigenin-7-oglucoside and luteolin, the flavonols quercetin and rutin and a number of phenolic acids such as caffeic, ferrulic, gallic, 3-hydroxy-4-methoxy-cinamic, homovanillic, hippuric, coumaric, siringic, p-hydroxy-benzoic acid, protocatechuic acid (Supplemental Table 1).

**Figure 1.**
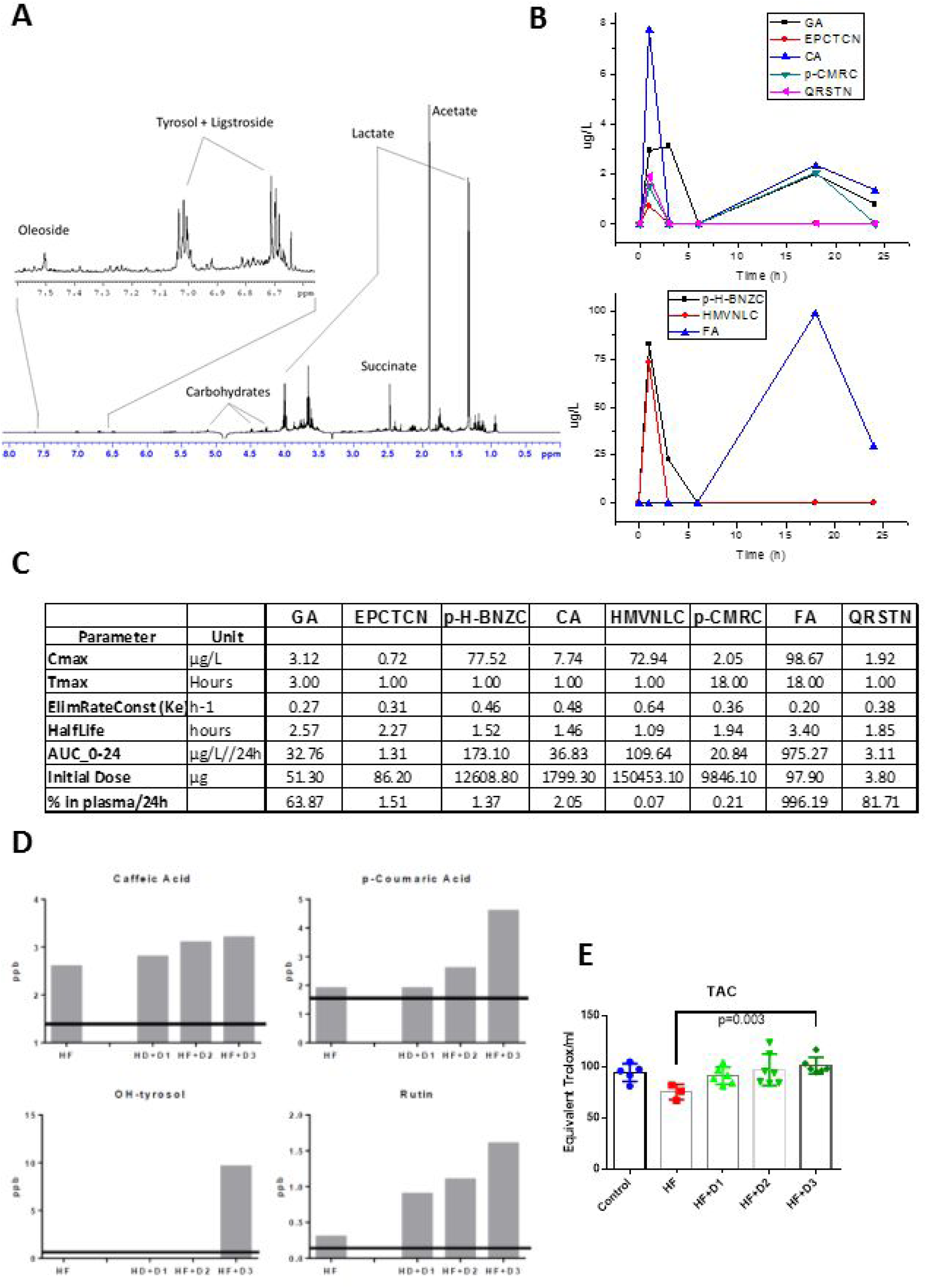
**A.** A characteristic ^1^H NMR spectrum of OLWPE in MeOD solvent and magnetic field 500MHz that shows its major constituents. **B.** The concentration (ppb:μg/L) of the different phenolic compounds detected in the plasma of rats by LC-ESC-MS/MS at different time points (1, 3, 6, 12, 18 or 24 h) after administration of OLWPE by gavage. Abbreviations: GA: Gallic acid, EPCTCN: Epicatechin, p-11-BNZC: p11-Benzoic acid, CA: Caffeic acid, HMVNLC: Homovanillic acid p-CMRC: p-Coumaric acid, FA: Ferulic acid, QRCTN: Quercetin **C.** Basic pharmacokinetic parameters of the phenolic compounds detected in OLWPE. Cmax: Maximum blood concentration, Tmax: Time required in order to achieve maximum blood concentration, AUC: Area under the curve “concentration vs time”, t_1/2_: half-life, Ke: Elimination rate constant. Initial Dose: the amount of each phenolic compound in OLWPE administered. % in plasma/24h: the percentage of each compound in OLWPE found in blood within 24h (% AUC/initial dose). Abbreviations: GA: Gallic acid, EPCTCN: Epicatechin, p-11-BNZC: p-11-Benzoic acid, CA: Caffeic acid, HMVNLC: Homovanillic acid p-CMRC: p-Coumaric acid, FA: Ferulic acid, QRCTN: Quercetin **D.** The concentration (ppb:μg/L) of the different phenolic compounds detected in the plasma of rats by LC-ESC-MS/MS after a 16-week consumption of the 3 different doses (D1-D3) of microencapsulated OLWPE extract in their food. *HF*: High fat diet. The parallel line to the x-axis represents the detection limit of the method for each compound. **E.** Total plasma antioxidant capacity (TAC) of animals fed either with normal diet (control) or a high fat diet (HF) with or without three different doses (D1, D2, D3) of microencapsulated OLWPE. Data are represented as mean ± SD.

The above extract was stabilized by micro-encapsulation and used in the subsequent long-term metabolic studies, described below.

### Animal study

#### OLWPE bioavailability and absence of toxicity in an animal model

Initially, in order to examine the bioavailability of the phenolic compounds in the extract, rats were given a single dose (corresponding to the dose D3 of the long-term study, see below and Material and Methods for further details) of OLWPE by gavage, blood was withdrawn at different time points (1-24h) and serum was analyzed by LC-ESI-MS/MS. As soon as one hour after treatment, a number of phenolic compounds have been detected in animal serum (Figure 1B and C) including epicatechin, quercetin, caffeic, gallic, coumaric, homovanillic, and p-hydroxy-benzoic acid. This early appearance of phenolics in the blood suggests an early gastric absorption. For caffeic, gallic, and coumaric acid, increased levels were also detected after 18 and/or 24h, indicating significant intestinal absorption, as well as, a possible increase as a result of (poly)phenol metabolism. As expected, oleuropein, was not detected since, due to its high molecular weight, it does not cross the intestinal barrier. Interestingly, its primary metabolite, hydroxytyrosol, was also not equally detected. Comparing the different AUC values, the compound with the greater bioavailability is ferrulic acid, followed by p-hydroxy-benzoic and homovanillic acid. The latter being a metabolite of hydroxytyrosol possibly explains its absence from the serum of treated animals.

Bioavailability data were also obtained during a long-term animal study, in which rats were fed with three different doses (D1<D2<D3) of microencapsulated OLWPE extract, for a period of 16 weeks. As shown in Figure 1D, increased levels of rutin, caffeic and p-coumaric acid, were detected in animal serum. These compounds were also present in rats fed only high fat food (HF), while their levels increased dose dependently when OLWPE was included in their diet. Moreover, in the highest extract dose (D3), hydroxytyrosol was detected, possibly as a product of long-term continuous oleuropein metabolism. Further evidence supporting the bioavailability of OLWPE were obtained by measuring the plasma total antioxidant capacity (TAC) of the 16 weeks-fed animals. TAC levels were dose-dependently increased in rats fed with different OLWPE extract; dose D3 exhibited a statistically significant increase compared to the high fat diet only group (Figure 1E).

Finally, at the end of the long-term animal study, rat livers and kidneys were examined in order to rule out any signs of toxicity that could be attributed to the polyphenolic extract. For this reason, organs were removed, formalin-fixed, paraffin embedded and sectioned for haemotoxylin-eosin staining. In Figure 2 such sections are presented for all study groups along with the levels of certain biochemical markers (GOT, GPT creatinine and urea,) of liver and kidney toxicity for all study groups. No signs of toxicity were identified, while a slight fatty liver infiltrates, as a consequence of high fat diet, were not modified by OLWPE.

**Figure 2.**
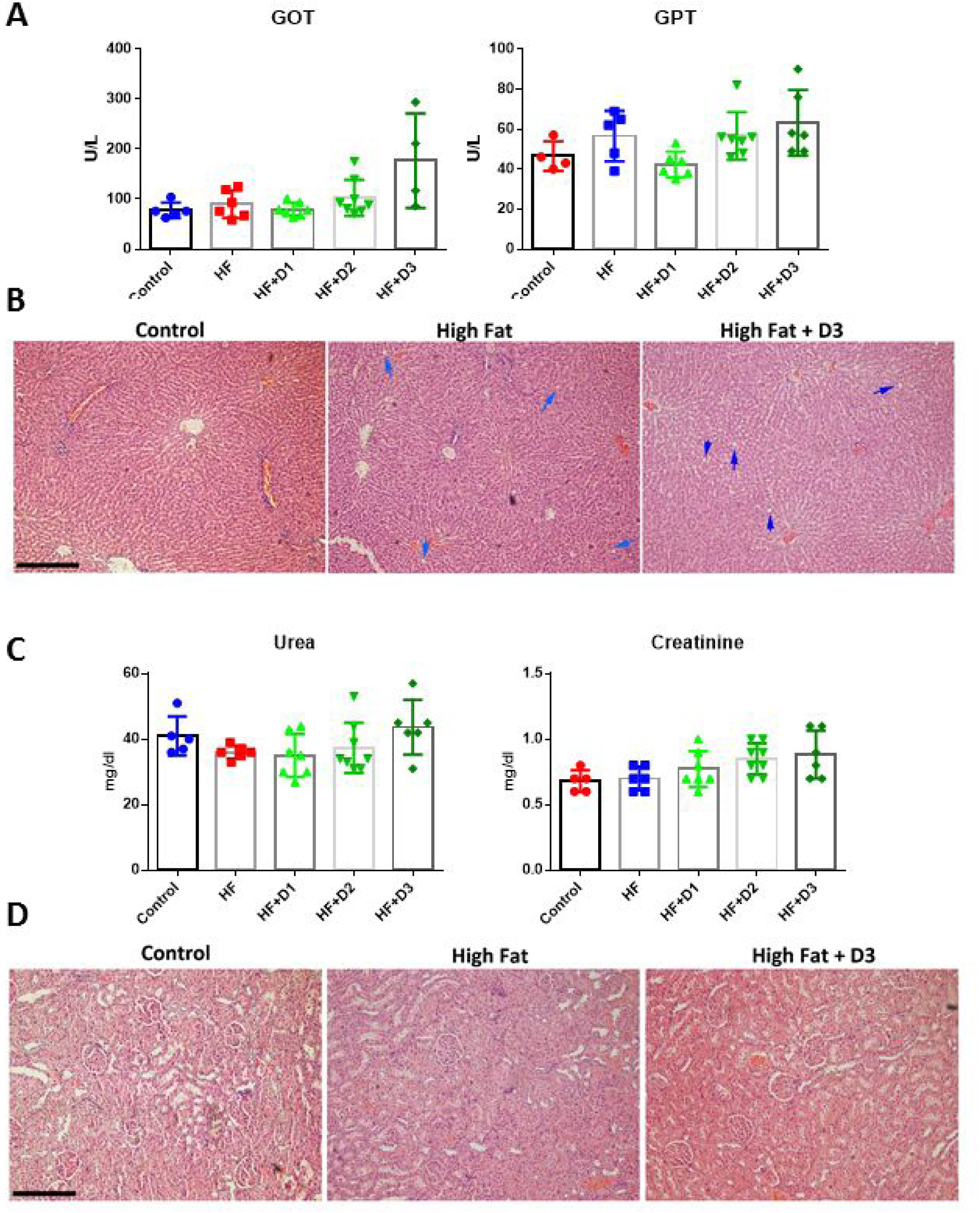
**A.** Changes in the concentration of the hepatic enzymes SGOT and SGPT in animals fed either with normal diet (control) or a high fat diet (HF) with or without three different doses (D1, D2, D3) of microencapsulated OLWPE for 16 weeks. Data are represented as mean ± SD. **B.** Representative microphotographs of hematoxylin-eosin stained liver sections of animals from the different study groups. Blue arrows indicate fat infiltration of the liver. **C.** Changes in the concentration of the urea and creatinine in animals fed either with normal diet (control) or a high fat diet (HF) with or without three different doses (D1, D2, D3) of microencapsulated OLWPE for 16 weeks. Data are represented as mean±SD. **D.** Representative microphotographs of hematoxylin-eosin stained kidneys sections of animals from the different study groups. Bars=100μΜ.

#### Effect of diets on rat weight

As it is shown in Figure 3A the final weight, as well as total weight gain, was elevated in rats receiving HF food alone or in combination with any of the tested OLWPE doses compared to the control rats. This indicates that HF diet induced obesity and OLWPE did not have any effect on body weight *per se*.

**Figure 3.**
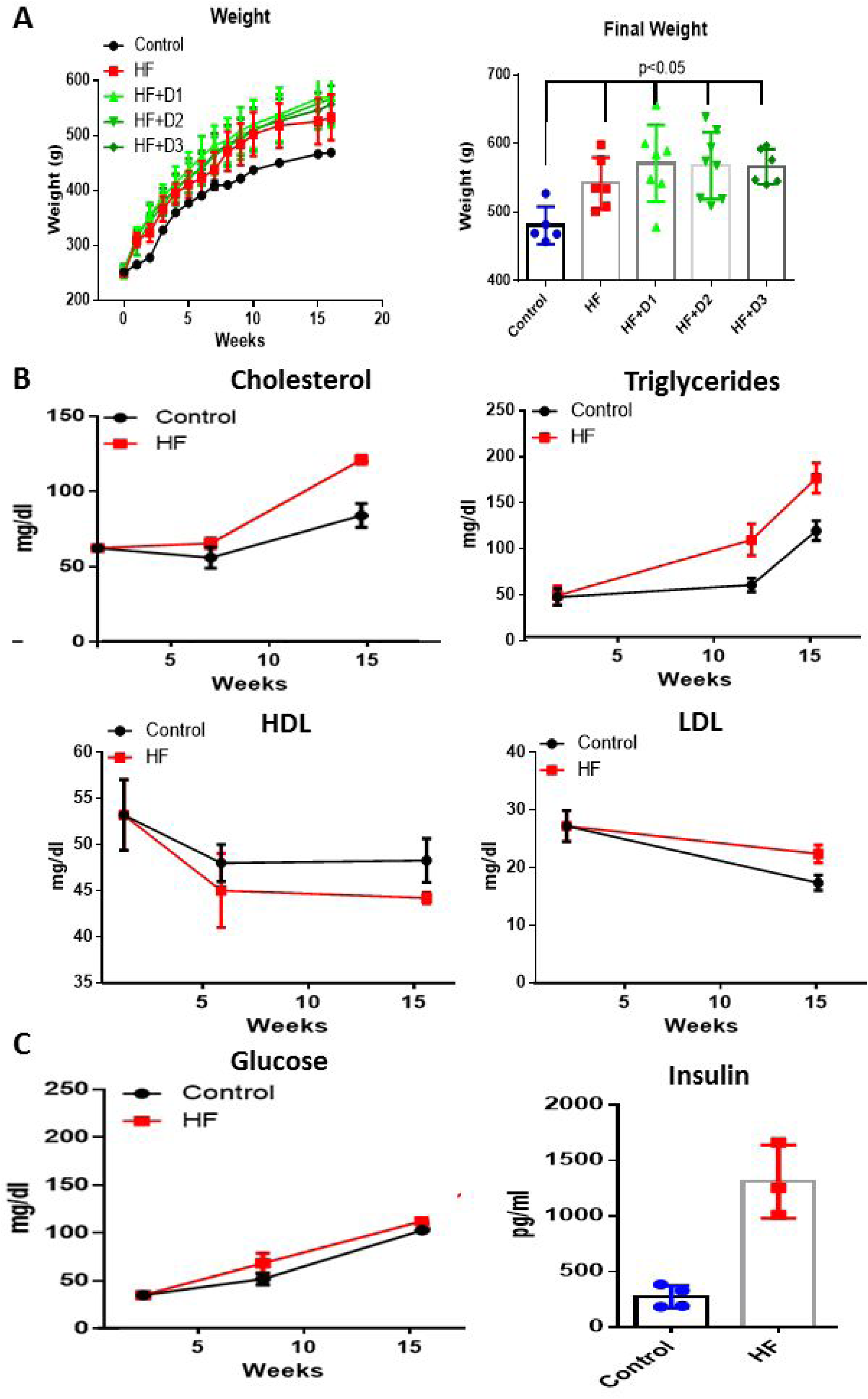
**A.** Changes in the weight of the rats fed either with normal diet (control) or high fat diet (HF) with or without three different doses (D1, D2, D3) of microencapsulated OLWPE within the period of 16 weeks. **B and C.** Changes of different metabolic parameters with time, in animals fed with high fat diet (HF) compared to normal diet (control). Data are represented as mean ± SD.

#### OLWPE significantly lowers circulating lipids and insulin levels

A sixteen-week high fat diet not only increased body weight but also induced additional features compatible with induction of metabolic syndrome: increased triglycerides, insulin and LDL and lower HDL (Figure 3B and D) were observed, while total cholesterol levels did not exhibit significant differences. Enrichment of HF diet with the different doses of OLWPE, significantly modified HDL and LDL levels (Figure 4). HDL levels were significantly elevated in rats receiving the highest (D3) dose, while LDL levels of all experimental groups receiving polyphenolic extract were significantly lower when compared to the HF only group. Accordingly, the HDL/LDL ratio (Figure 4B) was elevated in rats receiving doses 1 and 2, compared to rats of the high fat group. Additionally, insulin levels that were elevated in the HF only group compared to the control group, were decreased back to normal levels in rats receiving all three doses of the polyphenolic extract (Figure 4C). Finally, leptin that was significantly increased in HF diet rats was not modified by OLWPE (Figure 4D). This finding is in accordance with the lack of differences in the body weight between the HF only fed group and the groups with OLWPE in their diet. Similarly, no changes were observed in the levels of different proinflammatory cytokines like IL6 and TNFα (data not shown).

**Figure 4.**
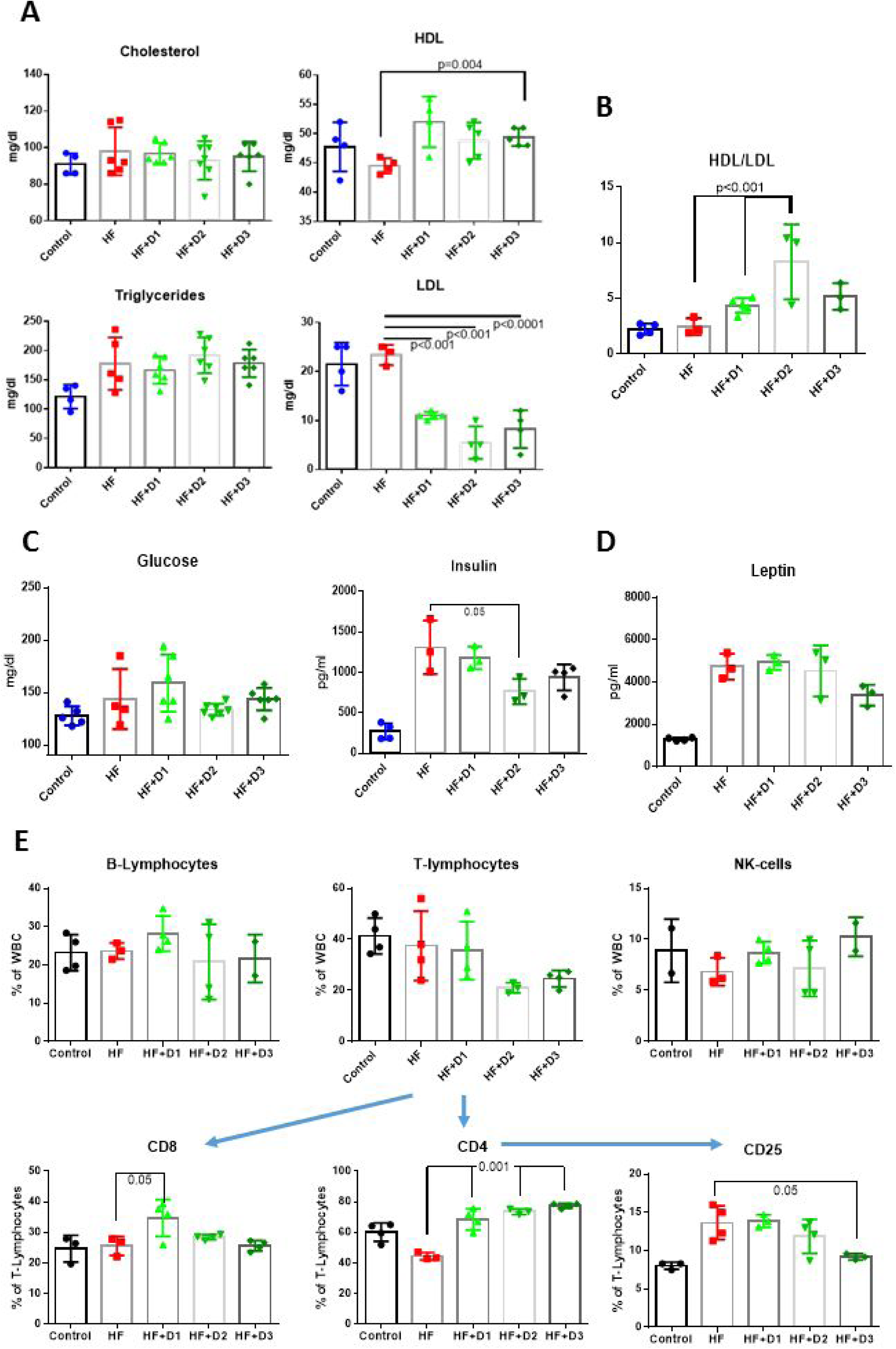
**A.** Changes in the lipidemic profile of animals fed either with a high fat diet (HF) with or without three different doses (D1, D2, D3) of microencapsulated OLWPE for 16 weeks. Values of the animals fed with a normal diet (control) are also presented. **B.** Changes of the HDL to LDL ratio at the different study groups. **C. and D.** Changes in glucose, insulin and leptin in animals fed either with a high fat diet (HF) with or without three different doses (D1, D2, D3) of microencapsulated OLWPE for 16 weeks. **E.** Immunophenotype: Changes in the different lymphocyte populations in animals fed either with a high fat diet (HF) with or without three different doses (D1, D2, D3) of microencapsulated OLWPE for 16 weeks. Values of the animals fed with a normal diet (control) are also presented. Data are represented as mean ± SD.

#### OLWPE decreases the levels of CD4^+^CD25^+^ T cells

Apart from the OLWPE-induced change of the aforementioned biochemical parameters, peripheral blood cell characteristics and immunophenotype were also examined. The major cell population numbers (red blood cells, lymphocytes, granulocytes and platelets) were not affected (Supplemental Figure 1). However, in an immunophenotype analysis (Figure 4E), CD4^+^ cells population was increased in an OLWPE dose dependent manner, while the CD4^+^CD25^+^ T regulatory cells were dose-dependently decreased by OLWPE and attained significant importance in the HF+D3 group, compared to the HF diet only group. All other immune populations did not present any significant differences among study groups.

### Human study

The above presented results from the animal study suggest that chronic consumption of OLWPE polyphenols, can improve the lipid profile of animals and reduce glucose levels and decrease insulin requirements. However, animal data are not easily extrapolated in humans, due mainly to a significantly different metabolism between the two species. Therefore, in order to explore whether the same effect can be obtained in humans, we have performed a proof of concept study, by administering microencapsulated OLWPE polyphenols (in a dose equivalent to the D2 dose used in the animal study, as the total polyphenol content included in the daily dose of olive oil approved by EFSA) in apparently healthy individuals. However, anthropometric data (Supplemental Table 2) showed that 19 out of the 35 participants were overweighted/obese (BMI >26) and 6 had a systolic blood pressure >130 mm Hg. The baseline biochemical analysis of our group revealed that 14 individuals had a fasting blood glucose >100 mg/dl, indicative of a pre-diabetic state, 26 presented a total cholesterol >200 mg/dl, 21 presented LDL cholesterol >130 mg/dl and 6 presented triglyceride levels >100 mg/dl. However, their HDL levels were astonishingly high (mean±SD 66.4±10.7 mg/dl), compatible with a high consumption of vegetables and olive oil, typical of a Cretan diet, followed by all participants.

#### NMR-based plasma metabolomics

At a first approach to identify changes in participants’ metabolism, when OLWPE is included in the diet of humans, serum samples from each participant before and after a four week OLWPE consumption were analyzed by NMR. Figure 5A depicts the OPLS-DA (Orthogonal Projection to Latent Structures – Discriminant Analysis) score plots before (ct1) and after (ct2) consumption of OLWPE, as obtained from the NMR metabolomic analysis of their serum lipids and water soluble metabolites (see Material and Methods for details). A clear separation of individuals is obtained from the OPLS-DA models, indicating that OLWPE consumption can be traced at both the water-soluble metabolite (R^2^X=0.80, Q^2^X=0.23) and serum lipid profile (R^2^X=0.91, Q^2^X=0.55), with the serum lipids model demonstrating a better discriminatory power (higher Q^2^X). In the case of serum lipids, buckets in the spectral region characteristic of LDL/HDL lipoprotein signal contribute significantly to the classification of individuals, indicating that OLWPE may affect their lipidemic profile.

**Figure 5.**
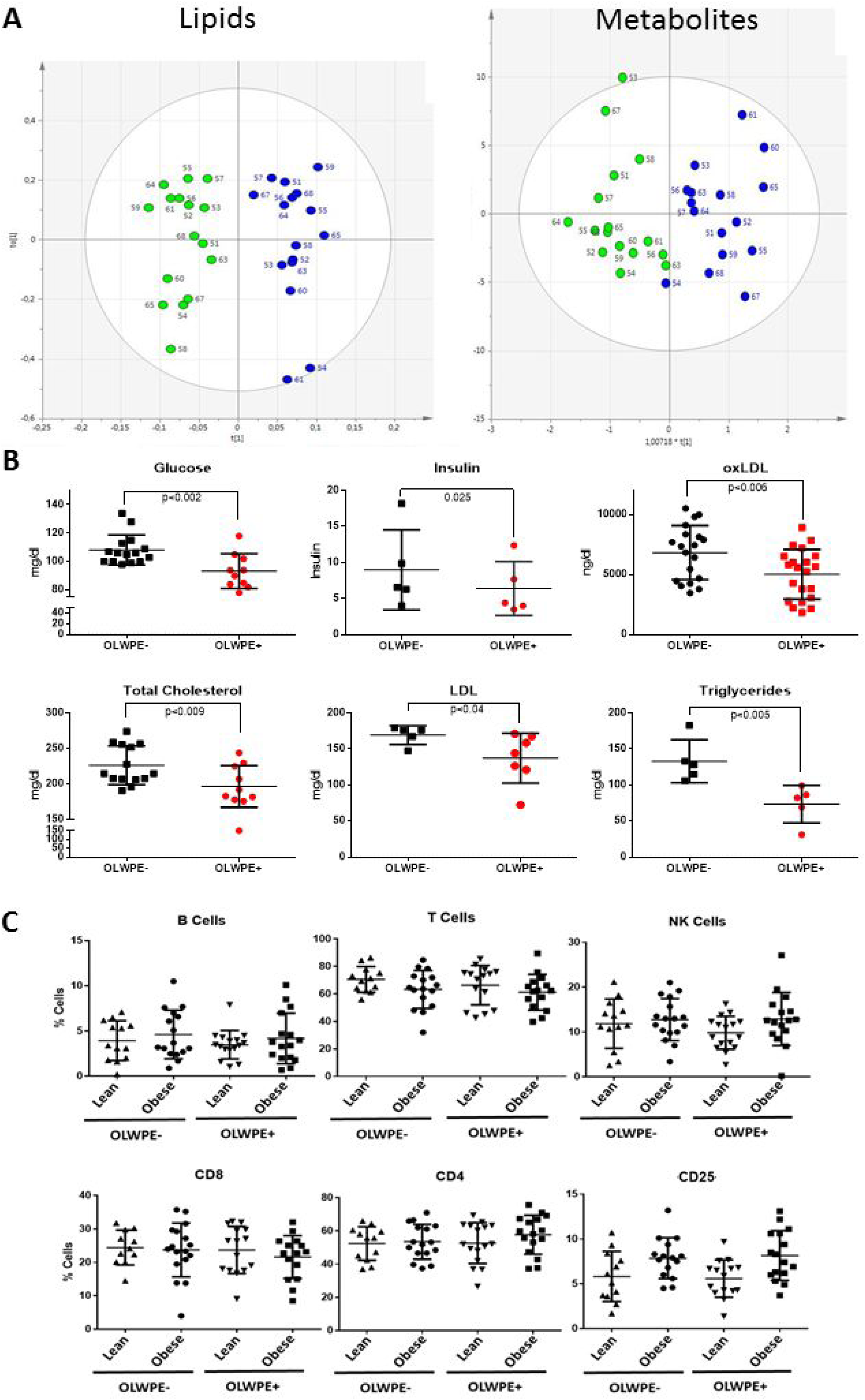
**A.** OPLS-DA models of ^1^H NMR spectroscopy obtained serum lipid and water soluble metabolite profiles of individuals (n=17) after consumption of a plain meat product (green) and one supplemented with OLWPE microencapsulated polyphenols (blue). **B.** The lipidemic and glycemic profile of individuals with at least two biochemical or anthropometric elements of cardio-metabolic risk (n=18) when consumed the meat product with the microencapsulated extract (OLWPE+) compared to their profile when the meat product was without the microencapsulated extract (OLWPE-). **C.** Immunophenotype: Changes in the different lymphocyte populations of individuals when consumed the meat product with the microencapsulated extract (OLWPE+) compared to their immunophenotype when the meat product was without the microencapsulated extract (OLWPE-). Individuals with BMI>26 (over weighted/obese) are presented as a separate group from the ones with BMI<26 (Lean). Data are represented as mean ± SD.

#### OLWPE decreases glucose and lipid levels

Based on the above metabolomic results and the beneficial effect of the microencapsulated OLWPE in animals, the lipidemic and glycemic profile was further explored. In normo-glycemic or normo-lipidemic individuals, the administration of the product did not modify blood glucose and lipid levels. However, in the sub-group of individuals with at least two biochemical or anthropometric elements of cardio-metabolic risk (n=18), we observed (Figure 5B) that, administration of OLWPE in a food matrix significantly reduced elevated glucose levels (p<0.002, n=15), while insulin levels were significantly reduced (p<0.03, n=6, paired analysis). In addition, OLWPE administration significantly reduced elevated total cholesterol (p<0.009, n=14), triglyceride (p<0.005, n=5) and LDL levels (p<0.01, n=5), while it decreased significantly oxLDL (p<0.01, n=18). oxLDL was also significantly reduced in the normolipidemic individuals, presumably as a result of the ingestion of polyphenols (paired t-test, p=0.0082, n=19) present in the extract.

Moreover, in the immunophenotype analysis (Figure 5C) no changes were observed in the major lymphocyte populations after OLWPE consumption. However, it should be noted that obese individuals have a slightly increased CD4^+^CD25^+^ T regulatory cells as expected from the animal study. Possibly the inability of OLWPE to decrease them can be attributed to the dose administered that corresponds to dose 2 of the animal model that equally has no significant effect on the different cell populations.

In all participants, no modification of circulating hepatic enzymes (SGOT, SGPT), urea or creatinine levels was observed (not shown), ensuring that this product does not express any hepatic or renal toxicity, at least for the period of its administration. Furthermore, no significant modification of body weight was observed, as expected, in the one-month interval of OLWPE consumption.

## DISCUSSION

Polyphenols (more than 8000 identified molecules containing a phenolic scaffold) constitute a large family of plant-derived compounds. They exhibit powerful antioxidant properties in parallel with a large number of biological actions, depending on their absorption and metabolism ^13,14^ A variable amount of ingested polyphenols can be found in blood ^15^, and can interfere with major cellular processes having a beneficial impact on cancer reviewed in ^13^, vascular function ^16,17^, and metabolism ^18,19^.

The beneficial role of olive oil consumption is now-a-days widely recognized, with the European Food Safety Authority (EFSA) approving two health claims regarding olive oil ^20^. They suggest its use to replace saturated fats in order to keep normal blood cholesterol levels and protect blood lipids from oxidative stress with the later effect to be achieved by olive oil polyphenols contained in a daily intake of 20 g of olive oil. In numerous studies, diets enriched either in virgin olive oil or following the pattern of the Mediterranean Diet (which is *per se* rich in olive oil, vegetables and legumes) have provided further evidence about the beneficial role of plant and olive oil antioxidant fractions in the prevention of cardiovascular disease ^21^, diabetes ^11,22,23^ and hyperlipidemia ^10^. Moreover, in an animal obesity and diabetes model, a polyphenol-enriched extract from olive leaves has been shown to reverse the chronic inflammation and oxidative stress that induces the cardiovascular, hepatic, and metabolic dysfunction ^12^. Olives are a rich source of polyphenols; during their harvesting and olive oil extraction, contained polyphenols are distributed between the oil and water phase, depending on the malaxation time and the applied temperature. Therefore, this water phase is a rich source of olive polyphenols, not yet exploited as a beneficial constituent of human functional foods/medicinal preparations. At this point It needs to be stressed that, according to a number of in vitro studies, the beneficial effect of food extracts is maximized when the total extract is used ^24,25^. This suggests: (1) either a synergistic effect of polyphenol molecules, that cannot be totally mimicked by the artificial combination of isolated phenolic molecules; or (2) that the effect of the extract is attributed to minor constituents, present in the total extract, beyond the lead molecules identified so far. Furthermore, an interesting observation that is in favor of the use of total plant extracts, is that isolated phenolic molecules with antioxidant properties (including vitamin C) in the context of a food matrix ^26^, may induce pro-oxidant activities, when administered isolated in vitro or in vivo ^27,28^.

In this light, in our study we used the total polyphenolic water extract from olives (OLWPE) that contains a number of different phenolic compounds (phenylethanoids, flavones, flavanols, flavonols and phenolic acids-See Results for details). Our findings can be summarized as follows: (1) We show that polyphenols and simple phenolic products are absorbed from different parts of the GI tract, as derived from acute metabolic studies; (2) The profile of absorbed polyphenols after chronic administration is different from that of an acute one; (3) Both in obesity-metabolic syndrome animal model and in humans, chronic administration of OLWPE in a food matrix results in improvement of glycemic control and lipid metabolic parameters, with no apparent toxicity. (4) In the animal model. OLWPE normalize the number of circulating CD4^+^CD25^+^ T regulatory cells that have been elevated in the metabolically dysfunctional high fat fed animals.

In our experimental conditions, animals fed with the high fat diet were significantly overweighed compared to those fed with a normal fat diet and exhibited metabolic dysfunction characterized by increased cholesterol, triglyceride, LDL and insulin levels, suggesting the establishment of insulin resistance. OLWPE ameliorates a number of these metabolic parameters such as lipidemic and glycemic profile, with lack of toxicity. These results are in accordance with a previous study on the effect of OMW biophenols on alloxan-induced diabetic rats ^29^. Especially in the case of increased LDL and decreased HDL levels, OLWPE seems to exhibit a highly beneficial effect. Additionally, even though fasting glucose was not significantly reduced, OLWPE reduced diet-induced hyperinsulinemia. Another interesting finding is that OLWPE can bring circulating CD4^+^ and CD4^+^CD25^+^ T regulatory cells that have been reduced and elevated respectively by high fat diet, back to normal levels. CD4^+^CD25^+^ T regulatory cells are important anti-inflammatory cells. However, their proportion in obesity-related metabolic disturbances is still controversial. There are studies reporting a reduction in obesity ^30-32^ and others, including our present findings, an increased number of peripheral blood Tregs ^33^. In fact, their percentage seems to be interdependent to their concentration in inflamed tissues. For instance inflamed obese visceral adipose tissue has been shown to have a reduced proportion of Tregs ^34,35^ and that could be a possible explanation for their increased number in the blood.

The above presented effects of OLWPE are greatly supported by our findings that the phenolic compounds of the extract are bioavailable (detected in rats’ sera) and the fact that the animals treated with the extract exhibited increased total plasma antioxidant capacity. Indeed several phenolic compounds present in the extract were detected in the blood of the animals (epicatechin, quercetin,caffeic, gallic, coumaric, homovanillic, and p-hydroxy-benzoic acid) as early as one hour after ingestion, pointing out an additional absorption via stomach that is followed by intestinal absorption ^36^. Although oleuropein was not detected, as expected, samples collected in longer time points after the OLWPE administration showed significant levels of its metabolite hydroxytyrosol. However, the aforementioned effects of OLWPE cannot be solely attributed to the detected molecules, due to certain limitations such as the doses used and the detection limit of the LC-ESCI-MS/MS method. Nevertheless, our data present a proof of the bioavailability of the polyphenolic olive extract, as previously described ^37^.

Obesity in humans is considered an emerging health problem ^38^ and in spite of the number of studies, its prevalence continues to rise. Diet is certainly very important to obesity incidence, the metabolic dysfunction that usually occurs in obese individuals and to its negative consequences, such as cancer ^39^, aging ^40^, cardiovascular disease ^41^ and a number of other pathologies ^42^. Metabolic dysfunction includes changes in their lipid and glucose metabolism, characterized by increased LDL cholesterol and low HDL levels, high glucose and insulin levels that can result to endothelial dysfunction, atherosclerosis and subsequent heart disease ^43^. It is therefore of great importance to find and exploit substances that will improve the above metabolic parameters. When healthy young individuals consumed OLWPE daily for a period of 4 weeks, metabolomic analysis revealed clear differences in their lipid and water soluble metabolites compared to the period that they did not consumed OLWPE.

Furthermore, a significant amelioration of specific metabolic parameters was observed in those individuals identified with elevated cardio-metabolic risk factors (at least 2 factors among fasting glucose and insulin, triglycerides, total and LDL cholesterol, n=18). These findings are in accordance to previous studies that report a beneficial effect of olive oil and its phenolic constituents on lipid profile ^44-49^. Moreover, oxidized LDL which is known to actively participate in atheromatous plaque formation and atherosclerosis, was significantly decreased in OMWPE-treated individuals, supporting the beneficial effect of OLWPE on cardiovascular risk factors. Additionally, it is of great importance that with the inclusion of OLWPE in the diet (both in animals and humans) a normal insulin sensitivity was restored. In fact, it is the first study reporting a direct effect of olive water extract polyphenols on fasting glucose and insulin levels, while all previous studies were conducted with olive oil see for example ^50, for a recent publication^. Finally, eventhough overweighted/obese individuals exhibited an increase in peripheral blood CD4^+^CD25^+^ T regulatory cells (as also observed in the animal model), OLWPE consumption had no effect mainly due to the given dose which was also ineffective in animals.

Overall, we conclude that OLWPE can exhibit a beneficial health effect, mainly by modifying circulating lipids and glucose/insulin levels, and most importantly without the presence of monounsaturated fats of olive oil, on which there are contradictory data concerning their role on development of insulin resistance, type 2 diabetes mellitus and impaired vascular integrity and cardiovascular disease ^51-54^. Under this scope we suggest that this microencapsulated polyphenolic extract could possibly be used in the development of “functional” OLWPE polyphenol-enriched foods or as an alternative/adjuvant therapeutic approach in hyperlipidemia and impaired glucose/insulin regulation.

## METHODS

### Isolation and characterization of OLWPE

Olive water total polyphenolic extract (OLWPE) was obtained by using olive mill waste water, immediately collected during olive oil production through centrifugation, passing through a multilevel ion-exchange proprietary resin filter (patent no. GR20030100295 20030708 & WO2005003037) and elution with ethanol (75%); ethanol was subsequently fully evaporated and the water extract was concentrated through a rotor evaporator. The total content of polyphenols was estimated by the Folin-Ciocalteu method ^55^ that has been modified in order to use small volumes. Briefly, 20 μl sample was mixed with 80 μl of distilled water, 400 μl Na2CO3 (10% Na2CO3 in 0.85 N NaOH), and 500 μl Folin-Ciocalteu reagent (10%). The mixture was allowed to stand for 1 hour in the dark and absorbance was measured at 750 nm. The total phenolic profile was expressed as Trolox (a water-soluble analog of Vitamin E) equivalents.

The specific composition of the polyphenolic content was obtained by using NMR spectroscopy and Mass Spectrometry. NMR spectroscopy experiments were conducted on a Bruker Avance III NMR spectrometer, operating at 500 MHz for the proton nucleus. OLWPE extracts’ NMR analysis was performed by mixing 100 μl of the sample with 400 μl MeOD. The mixture was vigorously shaken and then placed in a 5 mm NMR tube, where 1D (zg30, zgpr) and 2D dimensional (gCOSY, gHSQC, gHMBC) NMR spectra were acquired. ^1^H NMR spectra were acquired using the standard one-dimensional NOESY pulse sequence with water presaturation. Quantitative analysis was performed by the ChenomX software.

Mass spectrometric analysis was carried out on a ThermoFinnigan Liquid Chromatography/triple quadrupole mass spectrometer on Electrospray Ionization (LC–ESI-MS/MS). The experimental conditions for the mass spectrometric analysis were the following: negative ionization mode; capillary voltage 4000V; argon pressure 1mTorr. For quantification purposes data were collected in the selected ion monitoring (SIM) mode. The applicability and reliability of this analytical approach was confirmed by method validation and successful analysis of several samples with different matrices. Extraction of polyphenols from samples was performed using solid-phase extraction (SPE) Strata-X, 30mg/1mL (Phenomenex), a vacuum manifold, and a vacuum source.

### Plasma samples polyphenolic content analysis

Plasma samples were enzymatically hydrolyzed with β-glucuronidase/sulfatase from Helix pomatia (≥100,000 U/mL) at 37 °C for 45 min before polyphenol extraction; analysis was performed by SPE and LC–ESI-MS/MS respectively, as described above.

### Extract microencapsulation

OLWPE extract was microencapsulated using Maltodextrins and Spray Drying (Mean particle size 10μm) by XEDEV bvba (Zelzate, Belgium) in order to protect polyphenols from oxidation and heat, and simultaneously mask their unpleasant bitter taste.

### Animal Study

#### Short-term study

Male Sprague-Dawley rats (16 weeks old, weighting 400-500 gr), purchased from Harlan Laboratories were used (n=4). In each animal, a single dose of the extract (containing 3.42mg total polyphenols, corresponding to Dosage 3, see below) was given by gavage, directly to the stomach and the animals were single caged, had unlimited access to food and water and were kept under normal laboratory conditions. They were closely monitored for 24h and blood sample was collected at different time points (1, 3, 6, 18 and 24h), up to 24 hours. The specific polyphenolic content of their plasma at different time points was determined by LC-ESI-MS/MS under the experimental conditions described above. For each phenolic compound that has been detected the following pharmacokinetic parameters have been calculated: i) maximum concentration (Cmax), ii) time required to achieve maximum concentration (Tmax), iii) area under the curve (AUC), iv) half-life (t1/2), and v) elimination rate constant (Ke), according to ^56^ and using PK Functions for Microsoft Excel by Joel I. Usansky (http://www.boomer.org).

#### Long-term study

##### Diets

Normal food (2018S) and high fat (HF) food (TD.06414), containing 60% Kcal from fat were purchased from Teklad, Harlan Laboratories (Supplemental Table 3).

Both diets were acquired in their paste form, so that microencapsulated OLWPE could be better incorporated. OLWPE was given in three dosages: 0.375 mg of total polyphenols or 0.85 gr of microcapsules per Kg of food (Dosage 1, D1), 3.75 mg of total polyphenols or 8.5 gr of microcapsules per Kg of food (Dosage 2, D2) and 37.5 mg of total polyphenols or 85 gr of microcapsules per Kg of food (Dosage 3, D3). Dosage 2 corresponds to the total polyphenol content of 20g olive oil, being the daily dose of olive oil approved by EFSA that when used to replace saturated fat contributes to the maintenance of normal blood cholesterol levels and to the protection of blood lipids from oxidation ^20^.

##### Animals and experimental design

Male Sprague-Dawley rats (8 weeks old, weighting 230-270 gr), were purchased from Harlan Laboratories and used in our experiments. Animals were caged in groups of 3-4 rats, had unlimited access to food and water and were kept under normal laboratory conditions. They were randomly assigned in 5 study groups (n=8 animals per group): 1) the Control group (normal diet), 2) the high fat (HF) diet group, 3) the high fat + Dosage 1 (HF +D1) group, 4) the high fat + Dosage 2 (HF +D2) group and 5) the high fat + Dosage 3 (HF +D3) group. Rats were kept on these diets for 4 months *ad libitum*. During this period, animals were closely monitored, their weight was measured weekly. At the start and at the end of the study, animals were fasted for 12-14h and afterwards blood samples were collected, for complete blood cell counting, immunophenotyping and biochemical analysis. When animals were sacrificed at the end of the study, selected tissues (kidney, liver, lungs, fat and heart) were also kept for histological analysis.

Animal studies were approved by the School of Medicine, University of Crete Committee for animal welfare (Protocol no. 6174/7-5-14) and all experiments were performed in accordance with relevant guidelines and regulations.

##### Blood sample analysis

*<u>Complete blood cell count</u>* was performed at the University Hospital of Heraklion, Laboratory of Haematology, in an Abbott Emerald Hematology (Abbott, CA, USA) Analyzer according to standard operating procedures.

*<u>Immunophenotyping</u>* was performed as follows: 100μl of whole blood were incubated with the different fluorescently labelled antibodies for 30 min, followed by red blood cell lysis with the addition of 2 ml BD FACS Lysing solution (for a least 10 min) and afterwards each sample was analyzed in a flow cytometer (Attune^®^ Acoustic Flow cytometer, Applied Biosystems, Thermo Fisher Scientific, Waltham, MA USA) using lymphocyte population of at least 5000 cells. FITC mouse Anti-Rat CD4 (561834), PE mouse Anti-Rat CD45 (554878), PE mouse Anti-Rat CD8a (559976), PE mouse Anti-Rat CD25 (554866), FITC mouse Anti-Rat CD32 (550272) and FITC mouse Anti-Rat CD3 (561801) were from BD Pharmingen^®^ (BD Biosciences, San Jose, CA USA). FITC mouse Anti-Rat CD19 (MA5-16536) was from Thermo Fisher Scientific (Waltham, MA USA).

*Biochemical and Metabolic parameters*: All biochemical and metabolic parameters (glucose, triglycerides, total cholesterol, LDL cholesterol, HDL cholesterol, hsCRP, urea, creatinine, γGT, SGOT and SGPT) were measured according to standard operating procedures using Olympus AU2700 Analyzer at the Laboratory of Biochemistry of the University Hospital of Heraklion.

*Hormones and pro-inflammatory cytokines:* Insulin, leptin, TNFα, and IL6, were measured in duplicate using a multiplex kit (MILLIPLEX^®^ MAP Rat Adipokine Magnetic Bead Panel RADPKMAG-80K-04; Millipore Corp., St. Charles, Missouri,USA), in a Luminex 100 apparatus.

*Total Antioxidant Capacity (TAC)*: TAC was measured colorimetrically using the crocin bleaching assay as described in ^46^.

##### Histology

Toxicity was assessed using haemotoxylin-eosin staining in formalin-fixed paraffin-embedded sections of selected tissues (liver, kidney) for all study groups.

### Human study

#### Participants and study design

Thirty-five volunteers, without any diagnosed disease, participated in this study. The study was implemented in the following phases: participant recruitment using a number of exclusion criteria (Supplemental Table 4), baseline data collection, test phase I (4 weeks), washout period (2 weeks) and a test phase II (4 weeks). Participants were divided into two groups. Group A consumed the meat product containing the microencapsulated OLWPE in test phase I and the same meat product without the microencapsulated OLWPE in test phase II and group B in the reverse order. Blood sample collection and weight measurements were taken at the start of the study (baseline values, t=0) at the end of phase I (t=1) and at the end of phase II (t=2)

All participants gave written informed consent to participate in the study. The study was approved by the Ethics Committee of the Heraklion University Hospital (Protocol no 10714) and was performed in accordance with relevant guidelines and regulations.

#### Diet

All participants in each test phase were on a free diet (a 3 day food record at t0, which represents the participants’ usual diet before study, and at t1 and t2 was obtained. All participants adhered to Mediterranean diet as estimated by Mediterranean Diet Score^57^. One portion (30g) of a meat product with or without the microencapsulated OLWPE (containing 7mg Trolox equivalents of total polyphenols, that is the average amount that can be found in 20g of olive oil and corresponds to D2 of the animal study) was provided in each participant per day.

### Data collection

Different social - demographic data, such as date of birth, gender, citizenship, marital status, place of residence, profession and contact information were collected along with a number of Anthropometric measurements, including weight, height, waist and hip circumference and body mass index (BMI) was calculated. Additionally, blood pressure and pulse rate were monitored and several health habits, like alcohol consumption, smoking, individual’s medical history and the use of any medication were recorded. All patients were followed, once a week with telephone interviews and a complete physical examination at the beginning and the end of the corresponding period of intervention. At the end of each intervention period a complete biochemical and hematology workup was performed.

#### Blood sample analysis

##### Metabolomic analysis

The metabolomic analysis was performed according to published protocols by Beckonert et al. ^58^ and Dona et al. ^59^ Briefly, aliquots of human plasma (200 μl) were added in Eppendorf tubes to 400 μl of 0.9% saline solution, vortexed for 30 seconds, then centrifuged at 12,000g for 5 min at 4 °C and the sample (600 μl) was transferred into 5mm NMR glass tubes ^58^.

A Carr-Purcell-Meiboom-Gill spin echo sequence with presaturation was used for acquiring ^1^H NMR spectra and obtaining the low MW metabolite profile of plasma. A diffusion edited sequence (ledbpgppr2s1d) with bipolar gradients and LED scheme was used to suppress low MW compounds and obtain the ^1^H NMR spectrum of lipoproteins. ^59^ Low MW metabolites were quantified in the CPMG NMR spectra using the ChenomX suite (ChenomX, SA). Both CPMG and diffusion-edited LED spectra were bucketed and the data were used directly for metabolomics analysis, which was performed using the Simca 13.02 software package by Umetrics.

<u>Complete blood cell count,</u> the levels of different <u>biochemical and metabolic parameters</u> (glucose, insulin, triglycerides, total cholesterol, LDL cholesterol, HDL cholesterol, CRP, urea, creatinine, γGT, SGOT and SGPT) and <u>Total Antioxidant Capacity (TAC)</u> were obtained as described above.

<u>Immunophenotyping</u> was performed as described above using the following antihuman antibodies from BD (BD Biosciences, San Jose, CA USA): anti-CD45 PERCP(554878), anti CD3FITC (555332), anti-CD4 PE(555347), anti-CD8 PE (555635), anti-CD25 FITC (555432), anti-CD19 FITC (555412), anti-CD20 PE(555623), anti-CD16PE (555407), anti CD56 FITC (562794).

<u>Oxidized Low Density Lipoprotein</u> (OxLDL) was assayed using an ELISA kit (Cloud-Clone Corp. Houston, TX, USA), according to the manufacturer’s instructions.

### Statistical analysis

Statistical analysis was performed using SPSS, V21 (IBM Corporation, NY USA), Origin V8 (OriginLab Corporation, Northampton, USA) and Prism V6, (GraphPad Software, Inc La Jolla Inc).

## AKNOWLEDGEMENTS

The Authors would like to acknowledge *COOPERATION 2011* - Partnerships of Production and Research Institution in Focused Research and Technology Sectors – NRSF 2007-2013 (11ΣΥΝ_2_1785) for the financial support and would like to thank Olvos Science S.A. for its kind offer of the kit for assaying oxidized LDL.

## AUTHOR CONTRIBUTIONS

Conceived and designed the experiments: MK, EC, CL, Performed the experiments and analyzed the data: NK, VPA, EM, SK, EG, SK, MT, EM, NM, MN, EB, EGS and GN Participated in its design and coordination and helped to draft the manuscript: GN, EC, CL, AS, EGS Wrote the paper: MK, EC All authors read and approved the final manuscript.

## REFERENCES

1 Collaboration, N. C. D. R. F. Trends in adult body-mass index in 200 countries from 1975 to 2014: a pooled analysis of 1698 population-based measurement studies with 19.2 million participants. Lancet 387, 1377–1396, doi:10.1016/S0140-6736(16)30054-X (2016).

2 Despres, J. P. & Lemieux, I. Abdominal obesity and metabolic syndrome. Nature 444, 881–887, doi:10.1038/nature05488 (2006).

3 de Ferranti, S. & Mozaffarian, D. The perfect storm: obesity, adipocyte dysfunction, and metabolic consequences. Clinical chemistry 54, 945–955, doi:10.1373/clinchem.2007.100156 (2008).

4 Gregor, M. F. & Hotamisligil, G. S. Inflammatory mechanisms in obesity. Annual review of immunology 29, 415–445, doi:10.1146/annurev-immunol-031210-101322 (2011).

5 Despres, J. P. et al. Abdominal obesity and the metabolic syndrome: contribution to global cardiometabolic risk. Arteriosclerosis, thrombosis, and vascular biology 28, 1039–1049, doi:10.1161/ATVBAHA.107.159228 (2008).

6 Morange, P. E. & Alessi, M. C. Thrombosis in central obesity and metabolic syndrome: mechanisms and epidemiology. Thrombosis and haemostasis 110, 669–680, doi:10.1160/TH13-01-0075 (2013).

7 Alberti, K. G. et al. Harmonizing the metabolic syndrome: a joint interim statement of the International Diabetes Federation Task Force on Epidemiology and Prevention; National Heart, Lung, and Blood Institute; American Heart Association; World Heart Federation; International Atherosclerosis Society; and International Association for the Study of Obesity. Circulation 120, 1640–1645, doi:10.1161/CIRCULATIONAHA.109.192644 (2009).

8 Jick, H., Wilson, A., Wiggins, P. & Chamberlin, D. P. Comparison of prescription drug costs in the United States and the United Kingdom, Part 1: statins. Pharmacotherapy 32, 1–6, doi:10.1002/PHAR.1005 (2012).

9 O’Neill, S. & O’Driscoll, L. Metabolic syndrome: a closer look at the growing epidemic and its associated pathologies. Obesity reviews : an official journal of the International Association for the Study of Obesity 16, 1–12, doi:10.1111/obr.12229 (2015).

10 Damasceno, N. R. et al. Crossover study of diets enriched with virgin olive oil, walnuts or almonds. Effects on lipids and other cardiovascular risk markers. Nutrition, metabolism, and cardiovascular diseases : NMCD 21 **Suppl 1**, S14–20, doi:S0939-4753(10)00297-8 [pii]10.1016/j.numecd.2010.12.006 (2011).

11 Perez-Martinez, P., Garcia-Rios, A., Delgado-Lista, J., Perez-Jimenez, F. & Lopez-Miranda, J. Mediterranean diet rich in olive oil and obesity, metabolic syndrome and diabetes mellitus. Curr Pharm Des 17, 769–777, doi:BSP/CPD/E-Pub/000390 [pii] (2011).

12 Poudyal, H., Campbell, F. & Brown, L. Olive leaf extract attenuates cardiac, hepatic, and metabolic changes in high carbohydrate-, high fat-fed rats. The Journal of nutrition 140, 946–953 (2010).

13 Kampa, M., Nifli, A. P., Notas, G. & Castanas, E. Polyphenols and cancer cell growth. Rev Physiol Biochem Pharmacol 159, 79–113, doi:10.1007/112_2006_0702 (2007).

14 Spencer, J. P., Abd-el-Mohsen, M. M. & Rice-Evans, C. Cellular uptake and metabolism of flavonoids and their metabolites: implications for their bioactivity. Arch Biochem Biophys 423, 148–161 (2004).

15 Walle, T., Walle, U. K. & Halushka, P. V. Carbon dioxide is the major metabolite of quercetin in humans. The Journal of nutrition 131, 2648–2652 (2001).

16 Delmas, D. & Lin, H. Y. Role of membrane dynamics processes and exogenous molecules in cellular resveratrol uptake: Consequences in bioavailability and activities. Molecular nutrition & food research, doi:10.1002/mnfr.201100065 (2011).

17 Ghosh, D. & Scheepens, A. Vascular action of polyphenols. Molecular nutrition & food research 53, 322–331, doi:10.1002/mnfr.200800182 (2009).

18 Possemiers, S., Bolca, S., Verstraete, W. & Heyerick, A. The intestinal microbiome: a separate organ inside the body with the metabolic potential to influence the bioactivity of botanicals. Fitoterapia 82, 53–66, doi:S0367- 326X(10)00189-9 [pii]10.1016/j.fitote.2010.07.012 (2011).

19 Vetterli, L. & Maechler, P. Resveratrol-activated SIRT1 in liver and pancreatic beta-cells: a Janus head looking to the same direction of metabolic homeostasis. Aging (Albany NY) 3, 444–449, doi:100304 [pii] (2011).

20 EFSA Panel on Dietetic Products, N. a. A. N. Scientific Opinion on the substantiation of health claims related to olive oil and maintenance of normal blood LDL-cholesterol concentrations (ID 1316, 1332), maintenance of normal (fasting) blood concentrations of triglycerides (ID 1316, 1332), maintenance of normal blood HDL cholesterol concentrations (ID 1316, 1332) and maintenance of normal blood glucose concentrations (ID 4244) pursuant to Article 13(1) of Regulation (EC) No 1924/2006. EFSA Journal 9, doi:10.2903/j.efsa.2011.2044 (2011).

21 Nadtochiy, S. M. & Redman, E. K. Mediterranean diet and cardioprotection: The role of nitrite, polyunsaturated fatty acids, and polyphenols. Nutrition 27, 733–744, doi:S0899-9007(10)00406-5 [pii]10.1016/j.nut.2010.12.006 (2011).

22 Diez-Espino, J. et al. Adherence to the Mediterranean diet in patients with type 2 diabetes mellitus and HbA1c level. Ann Nutr Metab 58, 74–78, doi:000324718 [pii]10.1159/000324718 (2011).

23 Marin, C. et al. Mediterranean diet reduces endothelial damage and improves the regenerative capacity of endothelium. The American journal of clinical nutrition 93, 267–274, doi:ajcn.110.006866 [pii]10.3945/ajcn.110.006866 (2011).

24 Damianaki, A. et al. Potent inhibitory action of red wine polyphenols on human breast cancer cells. J Cell Biochem 78, 429–441, doi:10.1002/1097-4644(20000901)78:3<429::AID-JCB8>3.0.CO;2-M [pii] (2000).

25 Kampa, M. et al. Wine antioxidant polyphenols inhibit the proliferation of human prostate cancer cell lines. Nutr Cancer 37, 223–233 (2000).

26 Barbaste, M. et al. Dietary antioxidants, peroxidation and cardiovascular risks. J Nutr Health Aging 6, 209–223 (2002).

27 Young, A. J. & Lowe, G. M. Antioxidant and prooxidant properties of carotenoids. Arch Biochem Biophys 385, 20–27, doi:S0003-9861(00)92149-0 [pii]10.1006/abbi.2000.2149 (2001).

28 Zhang, P. & Omaye, S. T. Antioxidant and prooxidant roles for beta-carotene, alpha-tocopherol and ascorbic acid in human lung cells. Toxicol In Vitro 15, 13–24, doi:S0887-2333(00)00054-0 [pii] (2001).

29 D’Angelo, S. et al. Pharmacokinetics and metabolism of hydroxytyrosol, a natural antioxidant from olive oil. Drug metabolism and disposition: the biological fate of chemicals 29, 1492–1498 (2001).

30 Luczynski, W. et al. Generation of functional T-regulatory cells in children with metabolic syndrome. Archivum immunologiae et therapiae experimentalis 60, 487–495, doi:10.1007/s00005-012-0198-6 (2012).

31 Wagner, N. M. et al. Circulating regulatory T cells are reduced in obesity and may identify subjects at increased metabolic and cardiovascular risk. Obesity 21, 461–468, doi:10.1002/oby.20087 (2013).

32 Yun, J. M., Jialal, I. & Devaraj, S. Effects of epigallocatechin gallate on regulatory T cell number and function in obese v. lean volunteers. The British journal of nutrition 103, 1771–1777, doi:10.1017/S000711451000005X (2010).

33 van der Weerd, K. et al. Morbidly obese human subjects have increased peripheral blood CD4+ T cells with skewing toward a Treg- and Th2-dominated phenotype. Diabetes 61, 401–408, doi:10.2337/db11-1065 (2012).

34 Deiuliis, J. et al. Visceral adipose inflammation in obesity is associated with critical alterations in tregulatory cell numbers. PloS one 6, e16376, doi:10.1371/journal.pone.0016376 (2011).

35 Winer, S. & Winer, D. A. The adaptive immune system as a fundamental regulator of adipose tissue inflammation and insulin resistance. Immunology and cell biology 90, 755–762, doi:10.1038/icb.2011.110 (2012).

36 Bischoff, S. C. Quercetin: potentials in the prevention and therapy of disease. Current opinion in clinical nutrition and metabolic care 11, 733–740 (2008).

37 Mevel, E. et al. Olive and grape seed extract prevents post-traumatic osteoarthritis damages and exhibits in vitro anti IL-1beta activities before and after oral consumption. Scientific reports 6, 33527, doi:10.1038/srep33527 (2016).

38 Acheson, K. J. Carbohydrate and weight control: where do we stand? Current opinion in clinical nutrition and metabolic care 7, 485–492 (2004).

39 Bianchini, F., Kaaks, R. & Vainio, H. Overweight, obesity, and cancer risk. The Lancet. Oncology 3, 565–574 (2002).

40 Hart, R. W. et al. Adaptive role of caloric intake on the degenerative disease processes. Toxicological sciences : an official journal of the Society of Toxicology 52, 3–12 (1999).

41 Diniz, Y. S. et al. Diets rich in saturated and polyunsaturated fatty acids: metabolic shifting and cardiac health. Nutrition 20, 230–234, doi:10.1016/j.nut.2003.10.012 (2004).

42 Rolandsson, O. et al. Prediction of diabetes with body mass index, oral glucose tolerance test and islet cell autoantibodies in a regional population. Journal of internal medicine 249, 279–288 (2001).

43 Choi, C. U. et al. Statins do not decrease small, dense low-density lipoprotein. Texas Heart Institute journal / from the Texas Heart Institute of St. Luke’s Episcopal Hospital, Texas Children’s Hospital 37 421–428 (2010).

44 Corona, G. et al. The fate of olive oil polyphenols in the gastrointestinal tract: implications of gastric and colonic microflora-dependent biotransformation. Free radical research 40, 647–658, doi:10.1080/10715760500373000 (2006).

45 Gradolatto, A., Canivenc-Lavier, M. C., Basly, J. P., Siess, M. H. & Teyssier, C. Metabolism of apigenin by rat liver phase I and phase ii enzymes and by isolated perfused rat liver. Drug metabolism and disposition: the biological fate of chemicals 32, 58–65, doi:10.1124/dmd.32.1.58 (2004).

46 Kampa, M. et al. A new automated method for the determination of the Total Antioxidant Capacity (TAC) of human plasma, based on the crocin bleaching assay. BMC clinical pathology 2, 3 (2002).

47 Ruano, J. et al. Phenolic content of virgin olive oil improves ischemic reactive hyperemia in hypercholesterolemic patients. Journal of the American College of Cardiology 46, 1864–1868, doi:10.1016/j.jacc.2005.06.078 (2005).

48 Venturini, D., Simao, A. N., Urbano, M. R. & Dichi, I. Effects of extra virgin olive oil and fish oil on lipid profile and oxidative stress in patients with metabolic syndrome. Nutrition 31, 834–840, doi:10.1016/j.nut.2014.12.016 (2015).

49 Vissers, M. N., Zock, P. L., Roodenburg, A. J., Leenen, R. & Katan, M. B. Olive oil phenols are absorbed in humans. The Journal of nutrition 132, 409–417 (2002).

50 Fernandez-Castillejo, S. et al. Polyphenol rich olive oils improve lipoprotein particle atherogenic ratios and subclasses profile: A randomized, crossover, controlled trial. Molecular nutrition & food research 60, 1544–1554, doi:10.1002/mnfr.201501068 (2016).

51 Gómez-Romero, M., García-Villalba, R., Carrasco-Pancorbo, A. & Fernández-Gutiérrez, A. in Olive Oil - Constituents, Quality, Health Properties and Bioconversions (ed D. Boskou) pp. 333–356 ( INTECH, 2012).

52 Keita, H., Ramirez-San Juan, E., Paniagua-Castro, N., Garduno-Siciliano, L. & Quevedo, L. The long-term ingestion of a diet high in extra virgin olive oil produces obesity and insulin resistance but protects endothelial function in rats: a preliminary study. Diabetology & metabolic syndrome 5, 53, doi:10.1186/1758-5996-5-53 (2013).

53 Manach, C., Mazur, A. & Scalbert, A. Polyphenols and prevention of cardiovascular diseases. Curr Opin Lipidol 16, 77–84, doi:00041433-200502000-00013 [pii] (2005).

54 Manach, C., Scalbert, A., Morand, C., Remesy, C. & Jimenez, L. Polyphenols: food sources and bioavailability. The American journal of clinical nutrition 79, 727–747 (2004).

55 Singleton, V. L., Orthofer, R. & Lamuela-Raventós, R. M. in Methods in Enzymology Vol. 299 (ed Science Direct) 152–178 (Academic Press, 1999).

56 Loucks, J., Yost, S. & Kaplan, B. An introduction to basic pharmacokinetics. Transplantation 99, 903–907, doi:10.1097/TP.0000000000000754 (2015).

57 Panagiotakos, D. B., Pitsavos, C. & Stefanadis, C. Dietary patterns: a Mediterranean diet score and its relation to clinical and biological markers of cardiovascular disease risk. Nutrition, metabolism, and cardiovascular diseases : NMCD 16, 559–568, doi:10.1016/j.numecd.2005.08.006 (2006).

58 Beckonert, O. et al. Metabolic profiling, metabolomic and metabonomic procedures for NMR spectroscopy of urine, plasma, serum and tissue extracts. Nature protocols 2, 2692–2703, doi:10.1038/nprot.2007.376 (2007).

59 Dona, A. C. et al. Precision high-throughput proton NMR spectroscopy of human urine, serum, and plasma for large-scale metabolic phenotyping. Analytical chemistry 86, 9887–9894, doi:10.1021/ac5025039 (2014).

